# The effect of performance contingent reward prospects flexibly adapts to more versus less specific task goals

**DOI:** 10.1101/2024.08.07.607018

**Authors:** Nathalie Liegel, Daniel Schneider, Edmund Wascher, Laura-Isabelle Klatt, Stefan Arnau

## Abstract

In some situations, e.g., when we expect to gain a reward in case of good performance, goal-driven top-down attention is particularly strong. Little is known about the task specificity of such increases of top-down attention due to environmental factors. To understand to what extent performance-contingent reward prospects can result in specific and unspecific changes in cognitive processing, we here investigate reward effects under different levels of task specification.

Thirty-two participants performed a visual or an auditory discrimination task cued by two consecutive visual stimuli: First, a reward cue indicated if good performance was rewarded. Second, a task cue announced either which of the two tasks would follow (precise cue) or that both tasks would follow equally likely (imprecise cue).

Reward and task cue preciseness both significantly improved performance. Moreover, the response time difference between precisely and imprecisely cued trials was significantly stronger in rewarded than in unrewarded trials. These effects were reflected in ERP slow wave amplitudes: Reward and preciseness both significantly enhanced the contingent negative variation (CNV) prior to the task stimulus. In an early CNV time interval, both factors also showed an interaction. A negative slow wave prior to the task cue was also significantly enhanced for rewarded trials. This effect correlated with the reward difference in response times.

These results indicate that reward prospects trigger task-specific changes in preparatory top-down attention which can flexibly adapt over time and across different task requirements. This highlights that a reward-induced increase of cognitive control can occur on different specificity levels.

## Introduction

Constant switches between generalization on the one hand and specification on the other make human cognitive processing effective and flexible. This is exemplified by various cases: Categories and chunks facilitate memory; concrete examples facilitate understanding; schemata formation and knowledge transfer facilitate social interaction; precise and concrete guidelines facilitate adaptations to complex situations. Dealing with different specificity levels is crucial for all kinds of cognitive processes. Top-down attentional control, i.e., the selection and prioritization of those representations and processes which are goal-relevant (Buschman & Kastner, 2015; Oberauer, 2019), is no exception in this regard, as goals might be more or less precisely specified (Mansouri et al., 2020). More and less precise goals might even be active at the same time: For instance, looking for food in a supermarket can include visual attention to all kinds of cheese with a special focus on Camembert.

As goals need to be constantly updated in order to flexibly adapt to changing environments (Ullsperger, 2017), the question arises if an *update* of task goals also happens on different specificity levels. In the supermarket example, the information that the supermarket is going to close in several minutes might increase general arousal, it might intensify the visual search for cheese in general - including Camembert - or it might specifically enhance the search for Camembert. In the present study, we therefore examine the specificity of task goal updates. Note that according to the above-mentioned definition of top-down control, the specificity of task goals determines the specificity of top-down attention relying on those task goals. Hence, whenever we mention the specificity of task goals in this paper, we also refer to the specificity of the underlying top-down attention processes and vice versa.

Goals might be updated content-wise because of additional or new information, as it is the case during task switching (Kiesel et al., 2010). In this case, the specificity of the update obviously entirely depends on the new goal content. However, environmental signals can also lead to an increase of the effectiveness of attentional control without actually changing the content of current goals (Egner, 2017). It has been proposed that such an increase of attentional control is based on a strengthening of task representations in prefrontal cortex. As a result, certain task goals and processing pathways are more clearly prioritized over others than it would be the case with less attentional control (Botvinick & Braver, 2015; Egner, 2017; Etzel et al., 2016; Miller & Cohen, 2001). Such an increase in attentional control might be triggered by feedback signals in response to actions or by environmental cues prior to an action (Botvinick & Braver, 2015; Ullsperger, 2017).

In the context of performance monitoring, it has been shown that control increases can be both, very general or more task-specific, depending on the environmental factor triggering the update and depending on the concrete situation. For example, an increase of attentional control is more likely to occur on a general level when it is triggered by errors and negative feedback than when it is triggered by conflict detection (Forster & Cho, 2014). Errors have been found to trigger general autonomous nervous system changes or a task-unspecific slowing of responses (Ullsperger, 2017), whereas conflict adaptation is more task-specific (Egner, 2017). However, the specificity level of an update due to conflict adaptation still varies depending on the exact task set and paradigm (Braem et al., 2014). As little is known about the task specificity of goal adaptations that are triggered by environmental cues prior to an action, in this paper, we want to have a closer look on the task specificity of such a preparatory influencing factor.

Performance contingent reward prospects, that is expecting a reward in case of good task performance, is thought to result in a strengthening of task goals, an enhancement of attentional resource allocation and in an improvement of task performance (Botvinick & Braver, 2015; Etzel et al., 2016; Krebs & Woldorff, 2017; Shenhav et al., 2013). Evidence for increased attentional control comes amongst others from neuroscientific research. It has been found that reward prospects increased neural correlates of attentional resource allocation, namely BOLD activity in fronto-parietal brain networks, as well as P3 amplitude and contingent negative variation (CNV) amplitude in the ERP (Botvinick & Braver, 2015; Krebs & Woldorff, 2017). Interestingly, reward prospects seem to mainly influence resource allocation in preparatory time intervals, indicating an increased reliance on proactive control (Braver, 2012; Fröber & Dreisbach, 2014; Frömer et al., 2021; Jimura et al., 2010; Sawaki et al., 2015). Therefore, manipulating reward prospects seem to be well suited to examine the task specificity of an enhancement of *preparatory* attentional control.

There is evidence that performance contingent reward expectation can affect top-down attention on a general rather than a specific level (Liegel et al., 2022): Our research group examined task prioritization in a dual task paradigm using EEG. In each trial, a cue indicated with 100% validity if subjects had to focus on either a working memory or a cued number classification task. The number classification task was in some trials an odd-even decision, in others a smaller / bigger five decision. A feedback system rewarded good performance in the relevant task stronger than good performance in the irrelevant task. Resource allocation towards the number classification task correlated with decreased alpha power during number classification task performance. To test if this effect was classification specific or general, we conducted a multivariate decoding analysis on alpha power regarding the two possible number classifications, for working memory task relevant and number classification task relevant trials separately. The type of the number classification (odd / even vs. smaller / bigger five) could be decoded with high accuracy, with no significant difference regarding task relevance. The pattern of results strongly suggests that alpha power modulation in this study targeted only the more general level – the resource allocation between working memory and number classification task – and not the “deeper”, more specific level of the resource allocation between smaller bigger 5 and odd / even number classification. This study indicates that a goal update due to performance-contingent feedback reward can target general representations and resource allocation processes. However, the paradigm was multileveled and rather complex, and it will be interesting to see, if more straight-forward reward effects can also target general resource allocation.

The present study examined – to our knowledge for the first time – the specificity of performance-contingent reward effects. To this end, we conducted an event-related potential study during which we applied a monetary reward manipulation in a cued task switch paradigm (see Figure 1). Participants performed alternately in visual and auditory discrimination tasks. A task cue provided more or less specific information regarding the upcoming task: It either indicated that the auditory task would follow, or that the visual task would follow or that both tasks are equally likely. By manipulating reward prospects on the one hand and task specification on the other hand, we aimed to understand how reward prospects influenced specific and unspecific task goals and related preparatory top-down attentional processing. Similar paradigms have been applied before without reward manipulation. In these studies, one typically observes that people perform better in case of specific task cues than in case of unspecific task cues (Finke et al., 2012; Mazaheri et al., 2014; Sokoliuk et al., 2019). Such a behavioral effect indicates that individuals indeed make use of the pre-task information and adopt different task preparation strategies in specific and unspecific trials. In case that we can replicate the behavioral specificity effect, the paradigm hence is well suited to understand how reward affects specific and unspecific task preparation.

**Figure 1.**
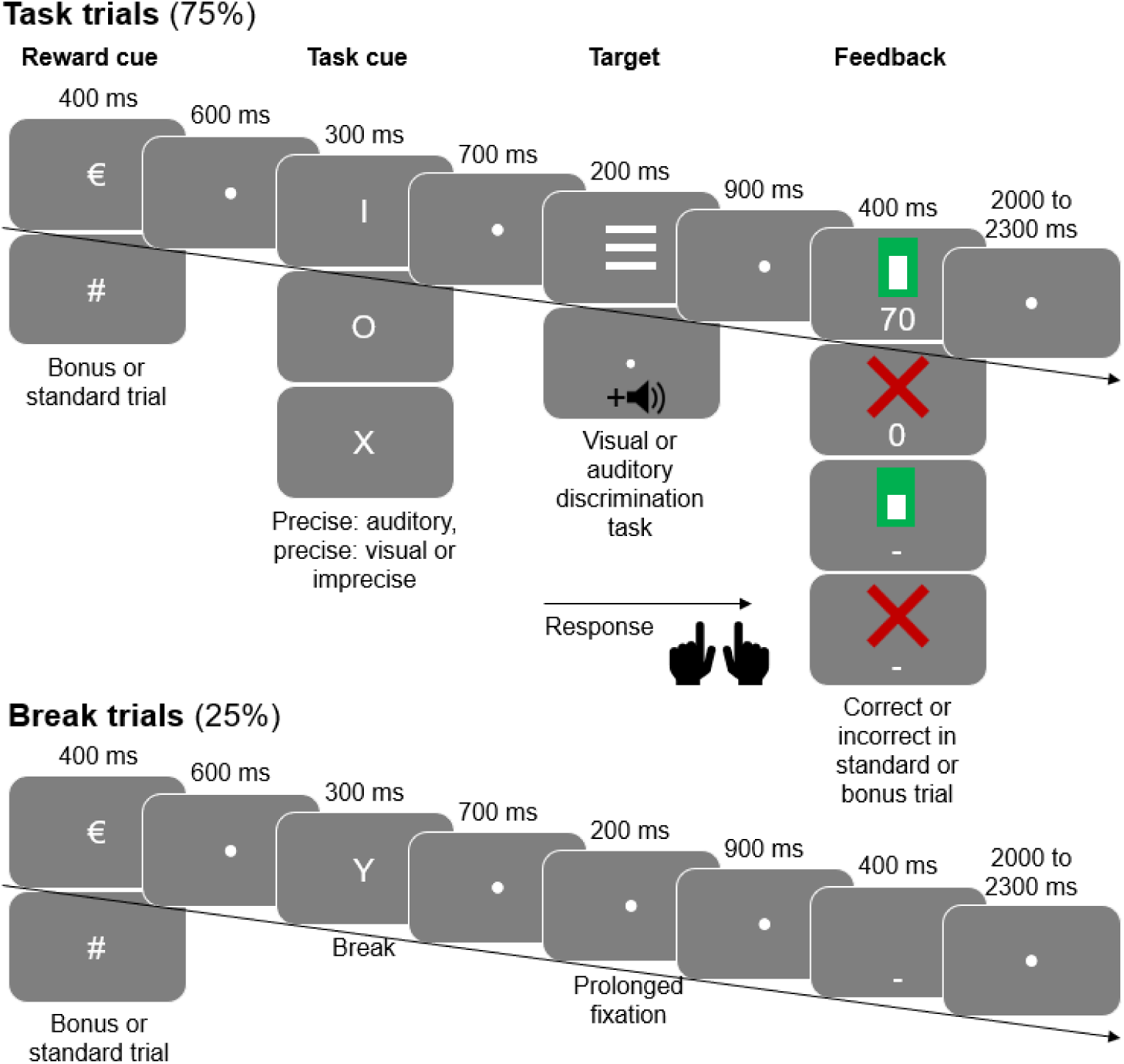
Experimental procedure. The majority of trials were task trials including either an auditory or a visual discrimination. A schematic illustration of those trial’s sequence is shown on top. ¼ of trials were break trials in which no task was done. Their sequence is depicted on bottom. Each trial started with a reward cue indicating if a monetary reward could be earned in case of good performance (bonus trials, cued by €) or not (standard trials, cued by #). Next, a task cue appeared – represented by a letter – indicating one of the following options: no task would follow (in the case of break trials), either an auditory or visual trial would follow with 100 % validity, respectively (precise task cue), either of the two tasks would follow with equal likelihood (imprecise task cue). The mapping of letters to task cue meaning was counterbalanced across participants. The task stimulus (target) was three horizontal or three vertical stripes in the case of a visual task or a pure tone with either a low or a high pitch in the case of the auditory task. In break trials, no task stimulus appeared. Participants classified the stripes’ orientation or the tone‘s pitch by pressing a button with either their left or their right hand. Finally, a feedback screen appeared consisting of an upper and lower part. In case of an incorrect or missing response, the upper part was a red cross. In case of a correct response the upper part was a white bar outlined by a green surrounding. The height of the white bar depended on response speed. In break trials, the upper part of the feedback screen was always empty. The lower part of the feedback screen was either a number (in case of task bonus trials) or a straight line (in case of break or standard trials). The number indicates how many points were earned in this trial. Points were translated to monetary incentive which was paid to the subjects after the experiment. (For illustrative purpose, stimuli are presented larger than in the actual task.).

Due to similar research in the context of performance monitoring and research on reward prospects in dual task scenarios, we hypothesized that the reward effect can happen on different specificity levels (Braem et al., 2014; Liegel et al., 2022; Ullsperger, 2017). We hence expected to find an effect of reward in case of both specific and unspecific task preparation in EEG and behavioral data. Beyond that, comparing both task specification levels, allowed us to examine if the reward effect differs for specific and unspecific task preparation. If that is the case, reward indeed affects task-specific processes in the case of precise task specification and task-unspecific processes in the case of imprecise specification. This indicates that the reward effect flexibly changes specificity levels dependent on the exact situation, as it is the case for conflict adaptation effects.

Finally, we also compared different time intervals during preparation to understand how reward specificity changes over time. Previous EEG studies manipulating performance contingent reward predominately used only one cue per trial: Either a cue indicated if a reward could be earned (Doñamayor et al., 2012; Fröber et al., 2021; Frömer et al., 2021; Kawasaki & Yamaguchi, 2013; Kostandyan et al., 2020; Plichta et al., 2013; Sawaki et al., 2015; Schneider et al., 2018; Seifert et al., 2006; van den Berg et al., 2014) or one cue coded both reward information and instructional content (Schevernels et al., 2014) or only a task cue containing non-reward related informative content about the upcoming task was applied and the reward information was not varied trial-by-trial (Arnau et al., 2023; Capa et al., 2013). Here, we used a reward cue and a task cue in order to divide the preparatory interval in a time interval of high uncertainty about the upcoming task and an interval with less uncertainty. The uncertainty was even bigger as our design included break trials in which no tasks had to be done: Those were randomly interspersed in between the normal task trials (see Figure 1). As a result, during the time interval prior to the task cue, participants did not know *if* a task would follow or not. Using this design, we aimed to understand if reward affects task goals before they are actually further specified in precisely cued trials; this would indicate that reward effects can vary in its specificity over time.

Taken together, in the current study, we aimed to compare effects of performance contingent reward prospects on more vs. less specific preparation processes while taking into account different temporal scales and task instructions.

## Methods

### Participants

Thirty-three volunteers participated in the study. The data of one participant was excluded from analysis due to bad behavioral performance (<75% correctly answered trials, 3 of 10 blocks under 70%, all other subjects >85% correct), resulting in a final sample of 32 participants (21 females) with a mean age of 24.9 years (*SD*=4.3). All participants had normal or corrected-to-normal vision, no psychological or neurological disorders and were right-handed according to the Edinburgh Handedness Inventory (Oldfield, 1971). As participants took part in an MRI scan after the EEG measurement, subjects with contraindications for MRI scanning were excluded. The study protocol was approved by the Ethical Committee of the Leibniz Research Centre for Working Environment and Human Factors. Informed consent was obtained from each participant and the study was conducted in line with the Declaration of Helsinki.

### Procedure

Participants performed a cued task switch paradigm framed by a monetary incentive delay task (see Figure 1). Each trial started with a reward cue indicating if a performance-contingent reward could be earned in the upcoming trial (bonus trials) or not (standard trials). Participants were instructed to always do the experimental task as quickly and accurately as possible but that they would only gain an extra incentive in bonus trials in case of good performance. It was stressed that in standard trials, they would not gain an extra incentive, independent of their performance. The reward cue was always a Euro-sign (€) for bonus trials and a hash mark (#) for standard trials. It was presented for 400 ms, followed by a fixation dot for 600 ms.

Next, a task cue appeared for 300 ms, followed by a fixation dot for 700 ms. The task cue was one of the four letters X, Y, I or O. Accordingly, the experiment comprised four different trial types that were mapped to a different task cue letter. The assignment was counterbalanced across participants. In precisely cued auditory trials, an auditory task would follow with 100% likelihood. Likewise, in a precisely cued visual trial, the task cue indicated with 100% validity that a visual task would follow. In trials with an imprecise task cue, an auditory or a visual task appeared with 50% likelihood, respectively. Finally, in break trials, the task cue indicated that no task would follow. Break trials were included in the experiment, although not analyzed, as we wanted to make sure that the task cue adds additional information and not only acts as an alerting signal in the case of the imprecise task cue. In this way, the time interval after imprecise task cues is comparable to the time interval after precise task cues as in both cases new information needs to be semantically processed and integrated into current goals.

Next, the target screen time interval followed for 200 ms. In break trials, no task was supposed to be done, therefore no target appeared and instead, the fixation dot remained on screen during that time interval. In precisely and imprecisely cued auditory trials, the task was to categorize a tone as having a high or low pitch, in precisely and imprecisely cued visual trials, the task was to categorize three parallel stripes as having either a horizontal or a vertical orientation. The response could be given upon target onset by pressing a button on a response device with either the left or the right hand. (The use of index fingers, thumbs or middle fingers were allowed as long as the same fingers were used for left and right-hand responses). Therefore, in visual trials, the target was one of two possible parallel stripes (one with a horizontal, one with a vertical orientation), in auditory trials, the target was one of two possible tones (either 1000 Hz for low pitch or 2000 Hz for high pitch). During the tone’s presentation in auditory trials, a fixation dot was presented on screen.

Finally, after another fixation dot, presented for 900 ms, a feedback screen was shown for 400 ms. Feedback was provided regarding response time, accuracy and performance-contingent reward (see also below). During the following inter-trial-interval, a fixation dot was again presented on screen. The inter-trial-interval’s duration was jittered from 2000 to 2300 ms in order to prevent systematic expectation effects during the baseline period prior to the reward cue.

During the experiment, participants sat in a comfortable armchair in a dimly lit room in a viewing distance of 1 m relative to a 32 in., 1920 × 1080 pixels VSG monitor (Display++ LCD) with 100 Hz refreshing rate. In ear headphones (Creative EP 630, Creative, Singapore) were worn. FreePascal software (https://www.freepascal.org/) was used as stimulus presentation software. Visual stimuli’s font type was Arial. Auditory pure tone stimuli were presented bi-laterally with an intensity of 65 dB. Throughout the experiment, the screen background color was grey (CIE1931: 0.287, 0.312, 10). Fixation dots (diameter: 0.18°), reward cues (height: 1°), task cues (height: 1°) and visual target stimuli (three parallel stripes, each stripe: 0.17° × 1.1°, distance between two stripes: 0.34°, respectively) were presented at the center of the screen in white (CIE1931: 0.287, 0.312, 50). The feedback screen consisted of an upper and a lower part: The upper part was empty in break trials. In task trials, it was either a red cross (CIE1931: 0.648, 0.341, 30) or a green bar (CIE1931: 0.306, 0.614, 30) with a white filling (CIE1931: 0.287, 0.312, 50), presented centrally. The red cross (height and width: 0.7°, line thickness: 0.1°) appeared in the case of an incorrect response or if the response time exceeded 1000ms relative to target onset (missing response). The green bar appeared in the case of correct responses and the height of its inner white bar depended on the preceding response time (outer green part: height: 1°, width: 0.5°, inner white part: width: 0.3°). The white bar had a minimum height of 0.2° in case of response times of 1000 ms and a maximum height of 0.83° in case of 100 ms or less. Response times between 100 and 1000 ms were accordingly assigned to bar heights in between, using a linear scale. The lower part of the feedback screen was presented in white (CIE1931: 0.287, 0.312, 50) at −1°. In standard and break trials, it was a straight line (height: 0.1°, width: 0.9°), in bonus trials, it was a number (height: 0.48°) representing the amount of feedback points earned in this trial.

Feedback points earned in bonus trials throughout the experiment were summed, translated to monetary incentive, and paid after the experiment. Each feedback point referred to €0.09 for the first 12 participants and to €0.14 € to the other participants. The relation was adapted due to a general increase of participant compensation in our lab because of an increase in minimum wage. In standard and bonus trials with breaks, no feedback points could be earned. In no break bonus trials, feedback points were zero if the response was incorrect or missing. In correctly answered bonus trials, feedback points (FP) were assigned to the response time in the current trial (RT) using the following formula:

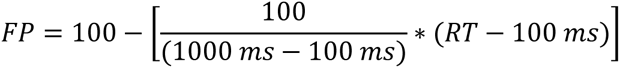

Participants were told that they could earn up to 100 feedback points per no break bonus trial and that the more feedback points they received the bigger their monetary incentive would be.

Each participant spent approximately 5.5 hours in the lab and received €30 as compensation for their time plus €22.10 to €48.90 extra incentive depending on their performance in the bonus trials (depending on feedback points). After the EEG measurement, an anatomical MRI measurement was done. The MRI data is not part of the present study. Prior to the start of the experiment, a 3D model of the electrode positions was obtained to align the MRI head shape to the EEG electrode positions using the Structure Sensor scanner (Occipital Inc., Boulder, Colorado). During that scan, the participant sat on a chair and relaxed (with eyes open or closed) for about 10 minutes. The experimental task was practiced step by step: First, each participant conducted 50 trials in which no reward cue was presented to get used to the cued task switch. Then, each participants conducted 40 - 120 additional practice trials based on the experimental trial sequence, this is, with reward cues. The large number of practice trials was required due to the complexity of the task.

In total, each participant performed 1280 trials, presented in randomized order, divided in 10 blocks of 128 trials each. At the end of each block, the participant was informed about the amount of additional monetary incentive due to feedback points, earned both in the respective last block and in all completed blocks. Each trial type (precisely auditory cued, precisely visual cued, imprecisely cued, break trials) appeared equally often. Half of the trials of each type were bonus trials, the other half standard trials. The levels of all experimental factors (bonus /standard, precise/imprecise, auditory/visual task, left/right response side) co-appeared equally often with the levels of each other experimental factor. In order to map the four task cue letters (I,O,X,Y) to a specific meaning, letters and meanings were grouped: I/O always appeared in one group and X/Y in the other. Task cue meanings were also grouped in precise task cues on the one hand and imprecise and break task cue on the other hand. For half of the participants, X/Y referred to precise task cues, for the other half to imprecise and break trial. This resulted in four different meaning-letter-associations (precise auditory, precise visual, imprecise, break = O,I,X,Y or I,O,Y,X or X,Y,O,I or Y,X,I,O). Each association co-occurred equally often with the four possible response side-response category mappings for visual and auditory task. The resulting 16 mapping versions were balanced across participants.

### EEG data acquisition and preprocessing

BrainVision Brainamp DC amplifier, BrainVision Recording software and 64 Ag/AgCl actiCAP slim active electrodes were used to record EEG data (BrainProducts, Gilching, Germany). The electrode montage was arranged according to the international 10–10 system with the ground electrode placed at AFz and the reference electrode at FCz. Data was recorded with a sampling rate of 1000 Hz and impedances were kept below 10 kΩ during recording.

EEG preprocessing was performed with EEGLAB (Delorme & Makeig, 2004) in combination with custom MATLAB code (R2021b, The MathWorks Inc., Natick, Massachusetts). Data were down-sampled to 500 Hz and high-pass filtered (FIR filter with Hamming window, order: 33001, transition band-width: 0.05 Hz, −6dB cutoff: 0.025 Hz, passband-edge: 0.05 Hz). Then, corrupted electrodes were identified by means of the built-in EEGLAb function *pop_rejchan* using kurtosis and probability criteria. Electrodes were rejected if their kurtosis exceeded +/-10 standard deviations of the mean based on the 20% trimmed kurtosis distribution of all channels. They were also rejected if their joint probability value exceeded +/-5 standard deviations of the mean based on a 20% trimmed joint probability distribution of all channels. Zero to six channels were removed per participant (*M* = 0.65 channels, *SD* = 1.38 channels). The function *pop_interp* was applied using spherical interpolation to restore previously removed channels; subsequently data were re-referenced to common average reference.

Starting here, data were preprocessed in two different ways:

First, in order to create optimal conditions to run an independent component analysis (ICA), data was down-sampled to 200 Hz, high-pass filtered a second time (FIR filter with Hamming window, order: 661, transition band-width: 1 Hz, −6dB cutoff: 0.5 Hz, passband-edge: 1 Hz) and low-pass filtered (FIR filter with Hamming window, order: 67, transition band-width: 10 Hz, −6dB cutoff: 45 Hz, passband-edge: 40 Hz). Baseline subtracted epochs ranging from −3600 to 5300 ms relative to reward cue onset were created with −200 to 0 ms relative to reward cue onset serving as baseline period. Epochs containing artifacts were automatically removed using probability criteria (EEG lab function *pop_autorej*). During this procedure, trials were rejected if they contained fluctuations larger than 1000 µV or if their joint probability exceeded +/-5 standard deviations of the mean based on a 20% trimmed joint probability distribution of all epochs of one channel. The joint probability was calculated in an iterative procedure and maximally 5% of trials per iteration were rejected. On average 5.19% of trials (*SD*=3.74) were removed. Then, an independent component analysis (ICA) was run on a data subsample (426 randomly drawn epochs, representing approximately 1/3 of all epochs). Prior to ICA, the rank of the data’s time point × channel matrix was determined to examine if there were potential linear dependencies amongst channels which could be detrimental to ICA calculation. If this was the case, a dimensionality reduction was done based on a standard PCA removing x dimensions that explained least variance, with x being 64 (number of all channels) minus the respective rank.

Second, in order to obtain preprocessed data for ERP analysis, ICA spheres and weights derived from the data set that was preprocessed for ICA were copied to the original data set, this is, to the 500 Hz-sampled data set obtained after re-referencing. Next, the EEGLab plugin ICLabel (Pion-Tonachini et al., 2019) was used to classify ICs. ICs that represented an artifact (eye movements /blinks, pulse or muscle artifacts, line noise, channel noise) with more than 50% likelihood according to ICLabel’s categorization were removed resulting in a mean rejection rate of 30.90% ICs (*SD* = 9.61%) per participant. Data was then divided in epochs ranging from −3600 to 5300 ms relative to reward cue onset. Again, the period from −200 to 0 ms relative to reward cue onset served as a baseline and the respective values were subtracted in each trial. Epochs containing amplitude fluctuations exceeding +/- 100 µV within the time interval of −500 to 3000 ms relative to the reward cue were rejected (*M*=3.17, *SD*=5.12). Data of epochs belonging to the same experimental condition and participant were then averaged in order to derive single-subject event-related potentials.

### Statistical analysis

To statistically analyze ERPs, we ran three cluster-based permutation tests (Maris & Oostenveld, 2007) by means of the Matlab toolbox Fieldtrip (Oostenveld et al., 2011). First, to test for a main effect of cue precision, precisely cued trials were compared to imprecisely cued trials. Second, to test for a main effect of reward, bonus trials were compared to standard trials. Third, to test for an interaction effect, the difference between bonus and standard trials was contrasted between precisely cued trials and imprecisely cued trials. The tests were computed over all 64 electrodes and across the whole epoch starting 200 ms prior to and ending 2700 ms after the reward cue. Note that break trials were not analyzed.

Test statistic for the permutation test were computed as follows: Two-sided paired t-tests were performed at each data point in electrode × time space: Sampling points with t-values exceeding a threshold of +/−2.0395 were identified and groups of more than two neighboring above-threshold points were defined as a cluster. Electrode neighborhood was specified using the Fieldtrip function *ft_prepare_neighbour* with default settings. The sum of all t-values within a cluster represented the cluster’s test statistic. To identify significant clusters, a non-parametric distribution of the test statistic was generated under the null hypothesis that there is no difference between the experimental conditions. Data representing the null hypothesis was created by randomly assigning experimental conditions to trials. The test statistic distribution was then calculated using a Montecarlo simulation with 4000 runs and applying the procedure described above in each run. Clusters in the real data were considered significant if their t-value sum exceeded the 97.5th percentile or fell below the 2.5th percentile of the H0 distribution.

For significant effects, adjusted partial eta squared effect size estimates (Mordkoff, 2019) were computed and reported by means of the symbol η.

## Results

### Behavioral data

Behavioral results are depicted in Figure 2. For response time analysis, a two-factorial repeated measures ANOVA was calculated on mean response times of correctly answered trials. Responses were significantly faster for bonus than for standard trials (main effect of reward, F(1,30) = 82.77, p < .001, η = .72) and faster in precisely cued than in imprecisely cued trials (main effect of preciseness, F(1,30) = 325.29, p < .001, η = .91). In addition, a significant interaction between both factors was found (*F*(1,30) = 11.01, *p* = .002, η = .24) with stronger differences between bonus and standard in precise than in imprecise task cue trials.

**Figure 2.**
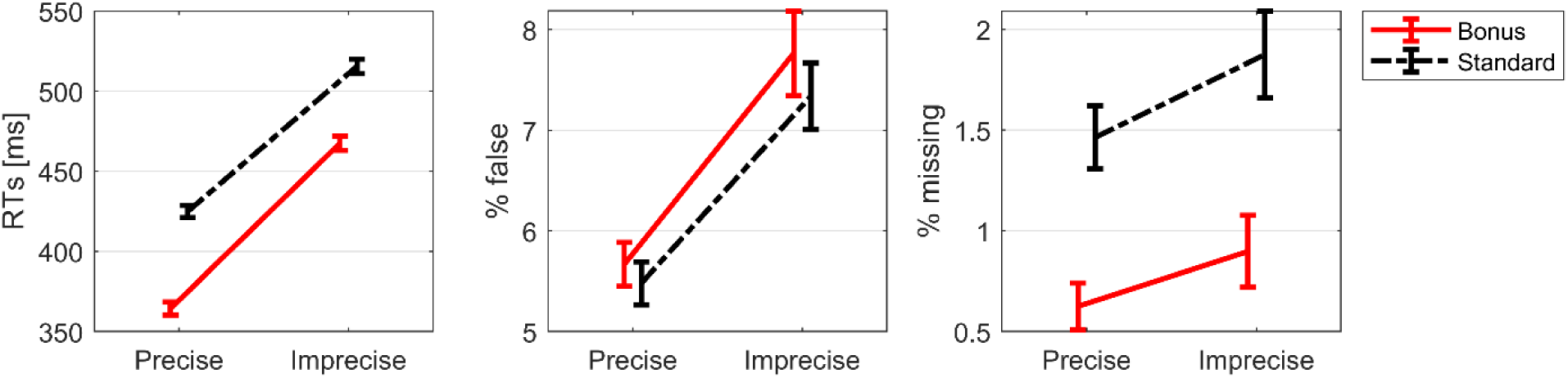
Behavioral results. Mean response times (left), error rates (middle), and omission error rates (right) for each experimental condition. Error bars represent within-subject standard errors of the mean (Cousineau-Morey method, for more information see Cousineau, 2005). Precise=trials with precise task cue; imprecise=trials with imprecise task cue, RTs=time between target onset and button press. A repeated measures ANOVA on error rates (% of incorrect answers) also revealed a main effect of the factor preciseness (*F*(1,30) = 22.18, *p* < .001, η = .40) with less errors in trials with precise compared to imprecise task cue. Neither the main effect of reward (*F*(1,30) = 0.85, *p* = .36), nor the interaction (*F*(1,30) = .19, *p* = .67) were significant. A third repeated measures ANOVA on omission errors, that is, the percent of trials in which no answer was given within 1000 ms after target onset, revealed only a significant main effect of reward (*F*(1,30) = 14.33, *p* < .001, η = 0.29). Neither the main effect of preciseness (*F*(1,30) = 4.34, *p* = .05), nor the interaction (*F*(1,30) = 0.17, *p* = .69) reached significance.

### Event-related potentials (ERPs)

Figure 3 shows ERPs at frontal, central and posterior electrodes for all experimental conditions. The figure shows clear differences between experimental conditions at fronto-central electrodes in preparatory time intervals. These slow wave differences start approximately 500 ms prior to the task cue and continue until the beginning of the target task. In line with Brunia and colleagues (2011), we label the slow wave in the time interval prior to the task cue *stimulus preceding negativity* (SPN) and the later slow wave that appears prior to the task stimulus *contingent negative variation* (CNV). Figure 3 also depicts condition differences in the amplitudes of the parietal P3 components (Polich, 2011) after the reward cue, the task cue and the task stimulus, as well as of the fronto-central N2 (Glazer et al., 2018) after the reward cue.

**Figure 3.**
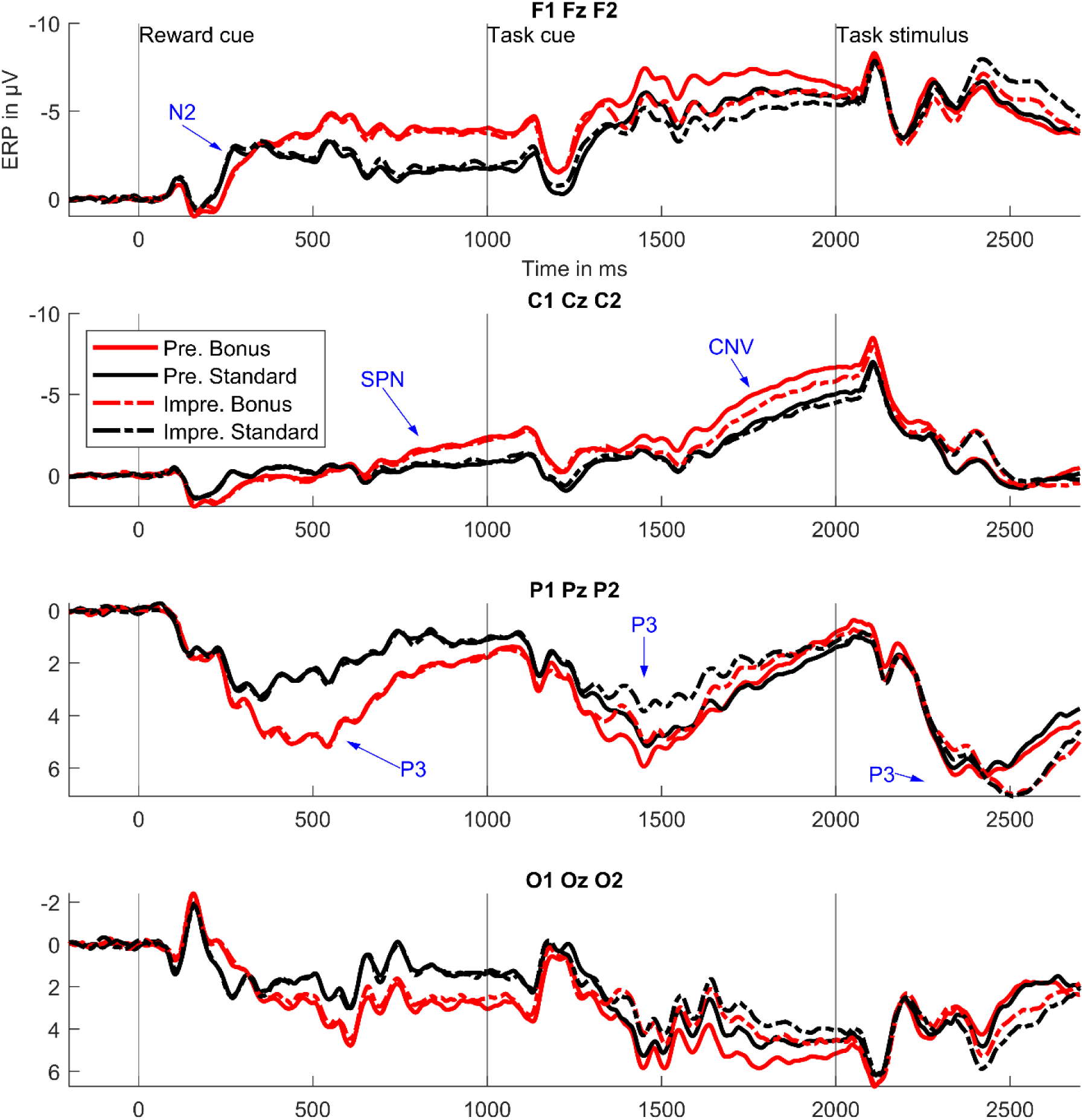
ERP time courses for different electrode clusters and experimental conditions. Pre.=precise task cue; Impre=imprecise task cue. ERPs were averaged over the three electrodes mentioned in the headline, e.g., over F1, F2 and Fz in the case of the top panel. Blue labeled arrows mark ERP components with strongest differences between experimental conditions (see Figure 4,6 and 9). CNV=contingent negative variation, SPN=stimulus-preceding negativity.

### Main effects of preciseness

To statistically analyze ERP data, we used cluster-based permutation tests including all electrodes and time points throughout the trial (−200 till 2700 ms relative to the reward cue). We chose this data-driven approach to describe effects comprehensively while also avoiding uncorrected multiple comparisons. A cluster-based permutation analysis testing for the main effects of preciseness revealed four significant clusters. Significant differences between precisely and imprecisely cued trials were found prior to the target as well as after the target. Cluster 1 (p<.001, precisely cued more negative than imprecisely cued) ranges from 1342-2016 ms, Cluster 2 (p<.001, precisely cued more negative than imprecisely cued) extends from 2374-2700 ms, Cluster 3 (p<.001, precisely cued more positive than imprecisely cued) encompasses time points from 1338-1904 ms. Finally, Cluster 4 (p<.001, precisely cued more positive than imprecisely cued) includes time points from 2248-2670 ms.

Figure 4 outlines those clusters in the electrode × time space and shows effect sizes at significant electrode – time combinations. Differences between precisely and imprecisely cued trials are most pronounced at around 1500 ms at frontal and parietal electrodes, at around 1700 ms at frontal electrodes and at around 2500 ms at parietal electrodes. Mapping those most pronounced differences to ERP components (see also Figure 3), the cluster-based permutation test revealed significant differences of preciseness on the amplitudes of the task cue P3, the target CNV and the target P3. Note that we use average-referenced data and hence component differences are mirrored to the other side of the scalp. Cluster 4 seems to represent such a mirror effect for the parietal P3.

**Figure 4.**
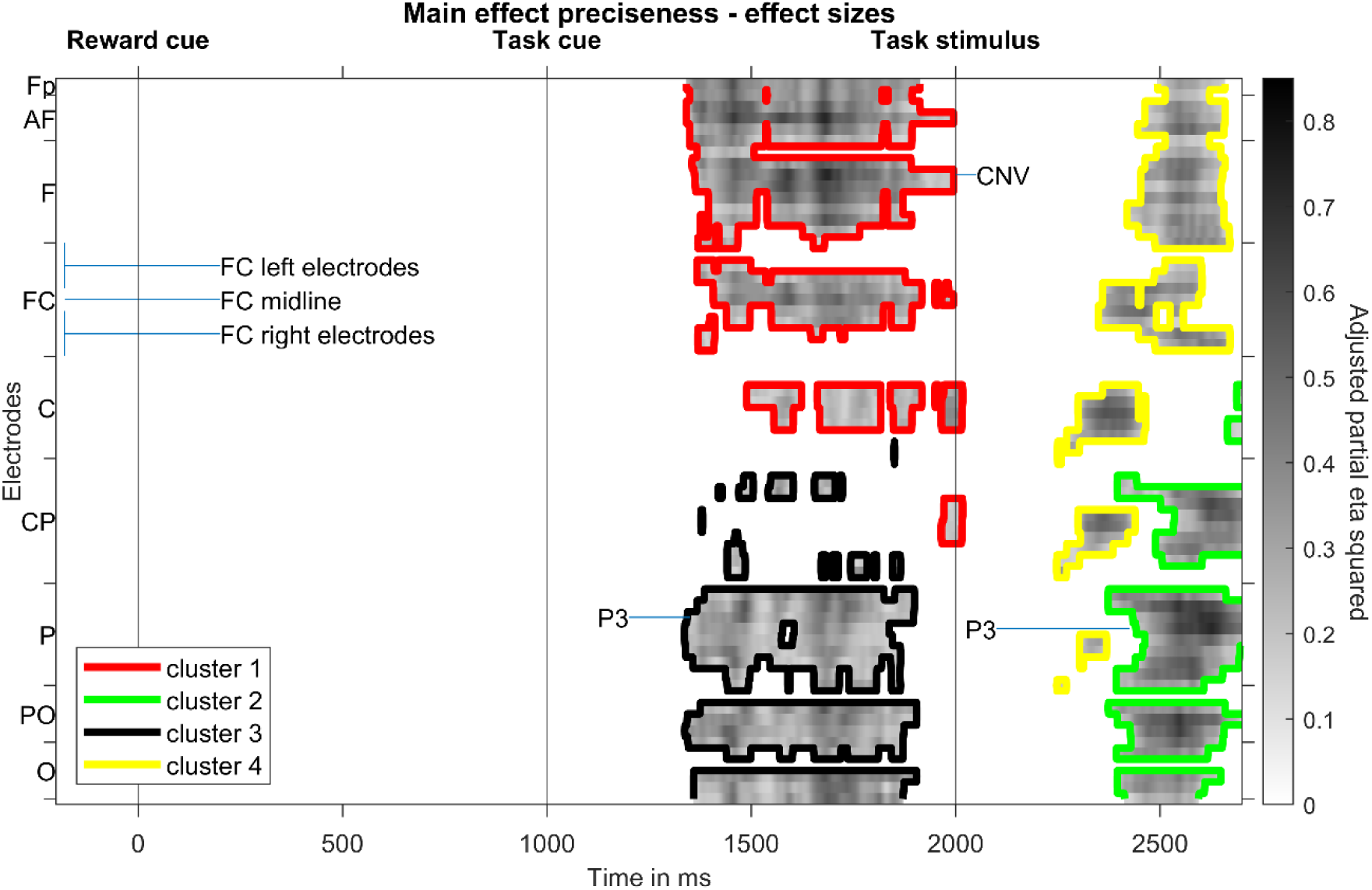
Result of the cluster-based permutation test on main effects of preciseness. Effect sizes (adjusted partial eta squared) for the precisely cued versus imprecisely cued comparison at significant time-electrode combinations are shown. Colored contour lines indicate to which cluster each time-electrode point belongs. Electrodes are lined up along the y axis, ranging from frontal to occipital rows, whereas each row’s electrodes are lined up from left to right (see labelling of FC electrode row); CNV=contingent negative variation.

Figure 5 shows topographic maps for time intervals during which significant differences between precisely cued and imprecisely cued conditions show strongest effect sizes. While the fronto-central target CNV and the parietal task cue P3 are bigger for precise task cues, the parietal target P3 is more pronounced for imprecisely cued task cues.

**Figure 5.**
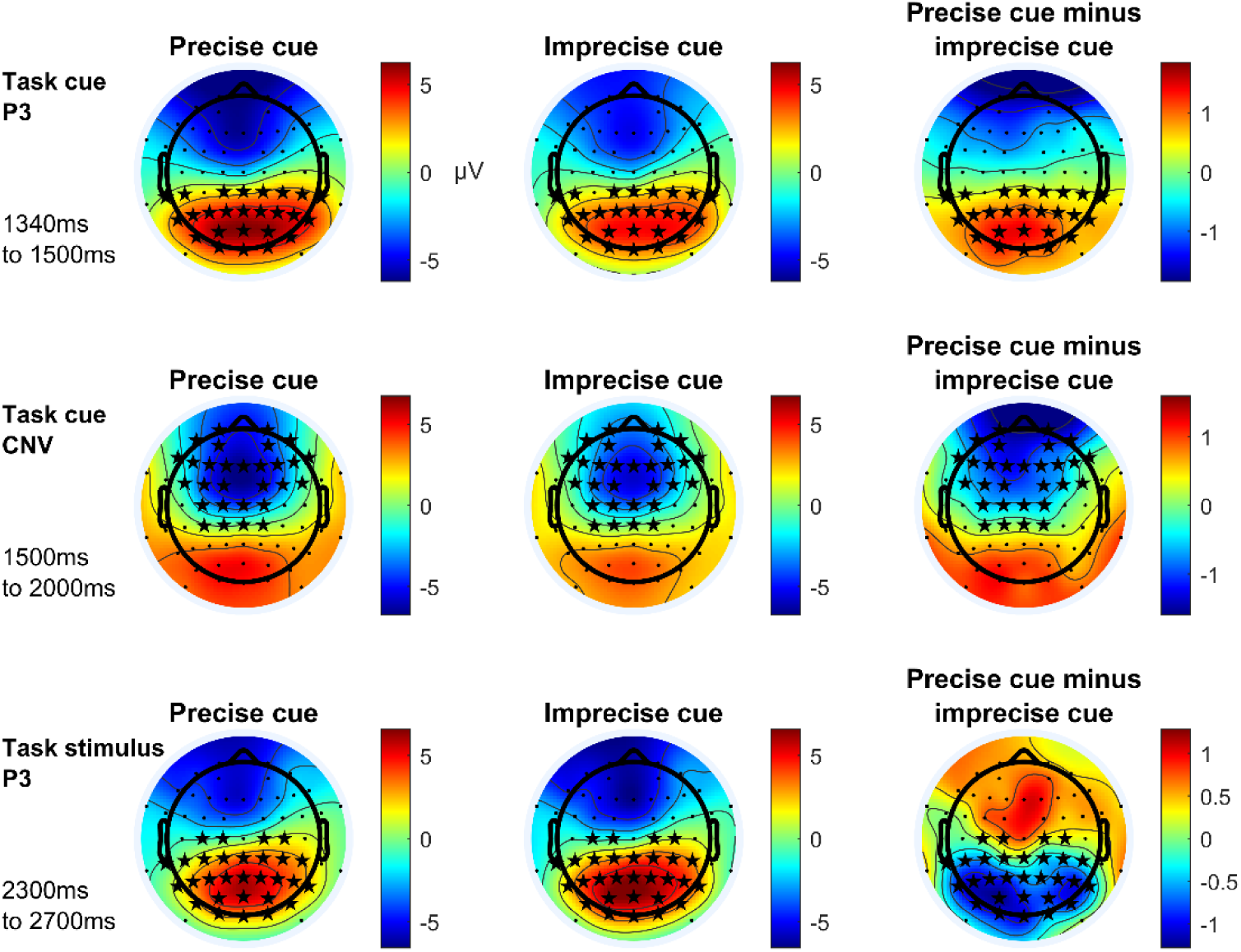
Topographic maps of ERP components with most pronounced effect sizes for main effects of preciseness. Depicted ERP components vary significantly, regarding their amplitude, between precisely and imprecisely cued trials according to a cluster-based permutation test. Asterisks mark electrodes showing significant differences at one or more time points of the respective time interval in the respective cluster. CNV=contingent negative variation. All times are given relative to the reward cue at 0 ms.

These results are in line with earlier reports on ERP data comparing precise and imprecise task cues (Finke et al., 2012).

### Main effects of bonus

A cluster-based permutation test comparing bonus and standard trials resulted in five significant clusters (see Figure 6). Cluster 1 (*p*<.001, bonus more negative than standard) comprises a time range from 352-1396 ms, Cluster 2 (*p*<.001, bonus more negative than standard) a time range from 1410-2184 ms, Cluster 3 (*p*=.02, bonus more negative than standard) a time range from 90-322 ms, Cluster 4 (*p*<.001, bonus more positive than standard) a time range from 196-1880 ms and Cluster 5 (*p*=.04, bonus more positive than standard) a time range from 92-184 ms.

**Figure 6.**
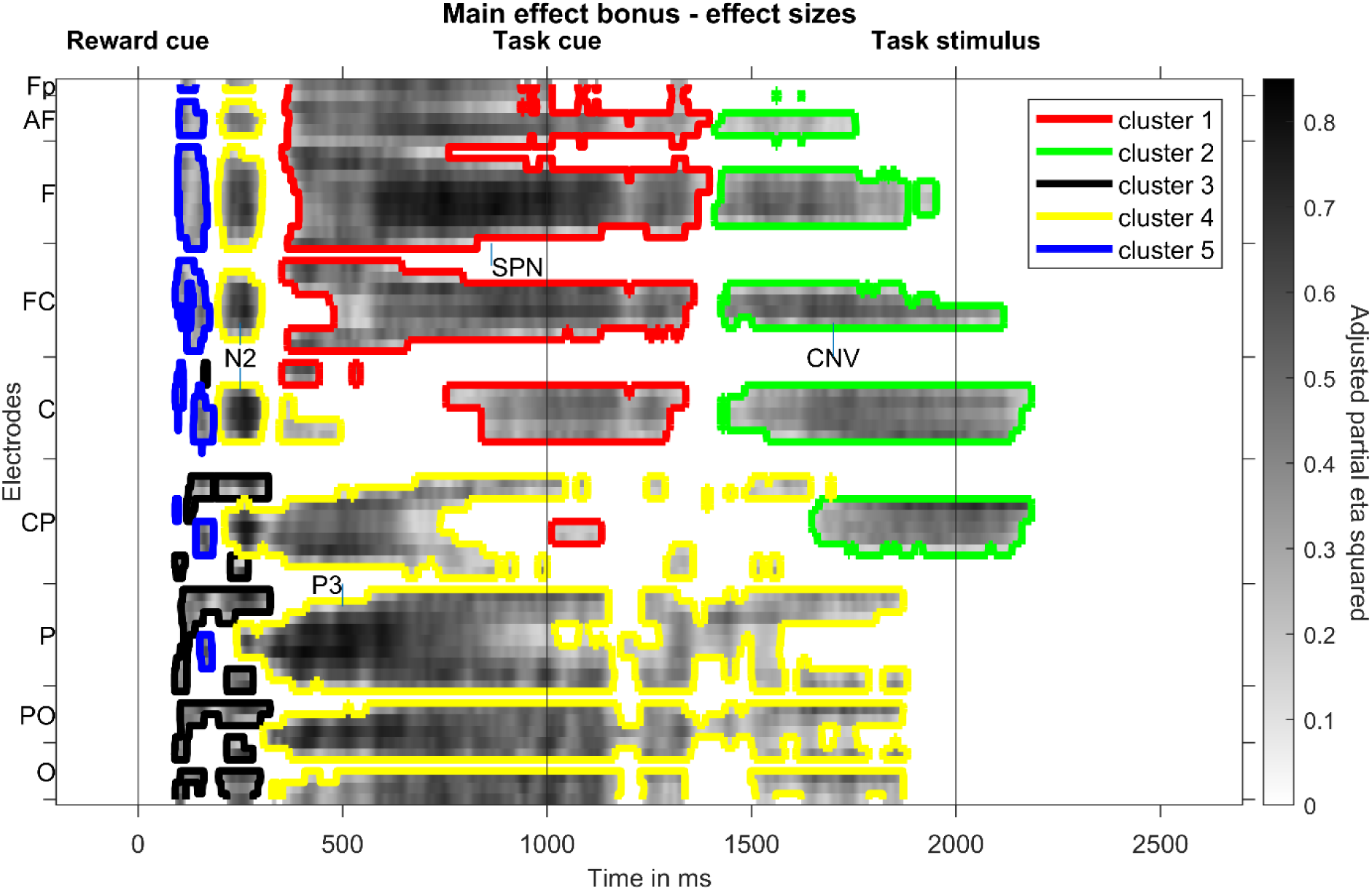
Result of the cluster-based permutation test on main effects of reward. Effect sizes (adjusted partial eta squared) for the bonus versus standard comparison at significant time-electrode combinations are shown. Colored contour lines indicate to which cluster each time-electrode point belongs. Electrodes are lined up along the y axis, ranging from frontal to occipital rows, whereas each row’s electrodes are lined up from left to right. CNV=contingent negative variation, SPN=stimulus-preceding negativity.

Effect sizes (see Figure 6) show strongest differences approximately 500 ms after the reward cue at parietal electrodes and in the 400 milliseconds preceding the task cue at frontal electrodes. In addition, strong differences can be found approximately 250 ms after the reward cue, as well as several hundred milliseconds prior to the target, at fronto-central electrodes, respectively. Mapping this to ERP components, reward expectation mainly modulated the reward cue N2, the reward cue P3, the task cue SPN and the task stimulus CNV.

Figure 7 shows topographic maps for those ERP components. Whereas frontocentral SPN and CNV slow wave amplitudes, as well as the parietal reward cue P3 amplitude is increased by reward expectation, the fronto-central N2 amplitude is decreased.

**Figure 7.**
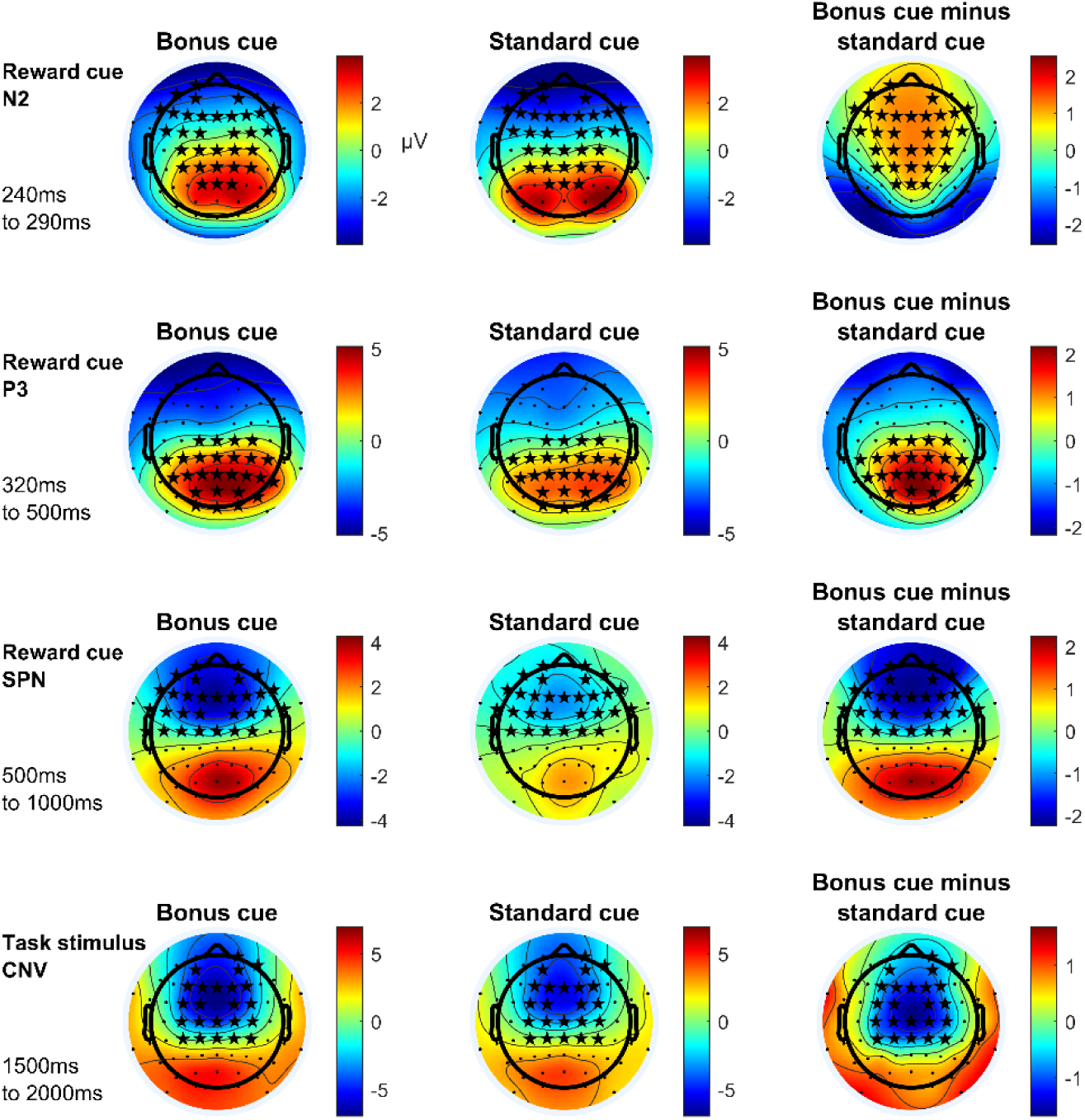
Topographic maps of ERP components with most pronounced effect sizes for main effects of bonus. Depicted ERP components vary significantly, regarding their amplitude, between bonus and standard trials according to a cluster-based permutation test. Asterisks mark electrodes showing significant differences at one or more time points of the respective time interval in the respective cluster. CNV=contingent negative variation, SPN=stimulus-preceding negativity. All times are given relative to the reward cue at 0 ms.

Like the analysis testing for the main effect of preciseness, the analysis testing for the main effects of reward resulted in plausible results: ERPs in response to reward cues have been investigated before and a stronger parietal P3b and a weaker central N2 amplitude have often been reported (Novak & Foti, 2015; Pornpattananangkul & Nusslock, 2015). Both has been interpreted as differences in cue evaluation, more specifically in stimulus categorization (P3b) and template mismatch evaluation (N2, for a review see Glazer et al., 2018). Likewise, the CNV prior to a task *stimulus* has earlier been reported to be increased by reward expectation (Glazer et al., 2018).

In contrast, a reward-related modulation of ERP activity prior to a task *cue* has not yet been investigated to our best knowledge. We therefore had a closer look at this effect. More exactly, we aimed to understand if the influence of reward on the SPN amplitude prior to the task cue reflects a behaviorally irrelevant by-product of reward cue processing or a functionally relevant preparation to the task. We calculated the Pearson correlation coefficient between the ERP bonus effect and the response time bonus effect across subjects. Hence, we subtracted the mean SPN amplitude for bonus trials from standard trials for each subject and used this difference for the correlation analysis. Time points and electrodes depicted in Figure 7 were used, i.e., the average for a time interval from 500-1000 ms relative to the reward cue at the electrodes Fp1, Fp2, AF7,AF3, AF4, AF8, F7, F5, F3, F1, Fz, F2, F4, F6, F8, FT7, FC5, FC3, FC1, FC2, FC4, FC6, FT8, T7, C5, C3, C1, Cz, C2, C4, CPz, CP2 was computed (see Figure 8). This analysis resulted in a significant correlation coefficient of *r*(30)=.59, *p*<.001, suggesting that the early, reward-related slow wave modulation prior to the task cue reflects task-related neural activity.

**Figure 8.**
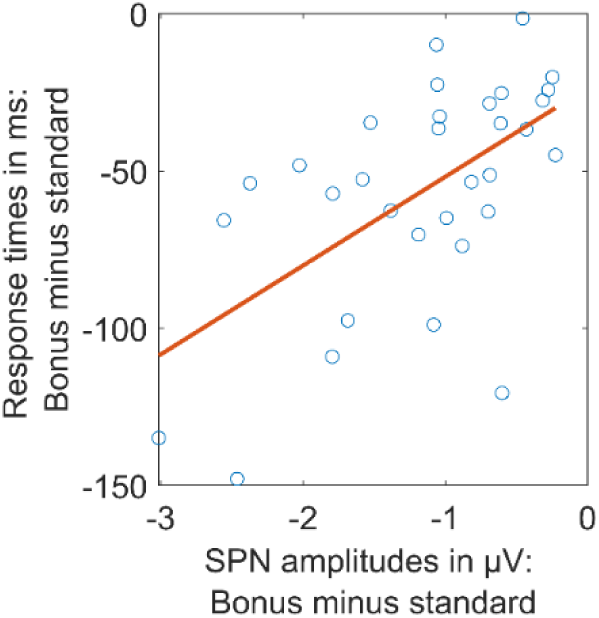
Brain-behavior correlation. Participants with strong response time bonus effects also showed strong ERP SPN bonus effects. Red line: Trendline, fitted according to least square principle.

### Interaction

As we were also interested how the bonus effect differs for precisely and imprecisely cued trials, we finally conducted a cluster-based permutation analysis testing for the interaction of both factors. To this end, the reward difference (bonus minus standard trials) of precise trials was compared with the reward difference of imprecise trials. A cluster (*p*=.009) of significant differences ranging from 1544-1634 ms was obtained with most pronounced effect sizes at central electrodes (see Figure 9).

**Figure 9.**
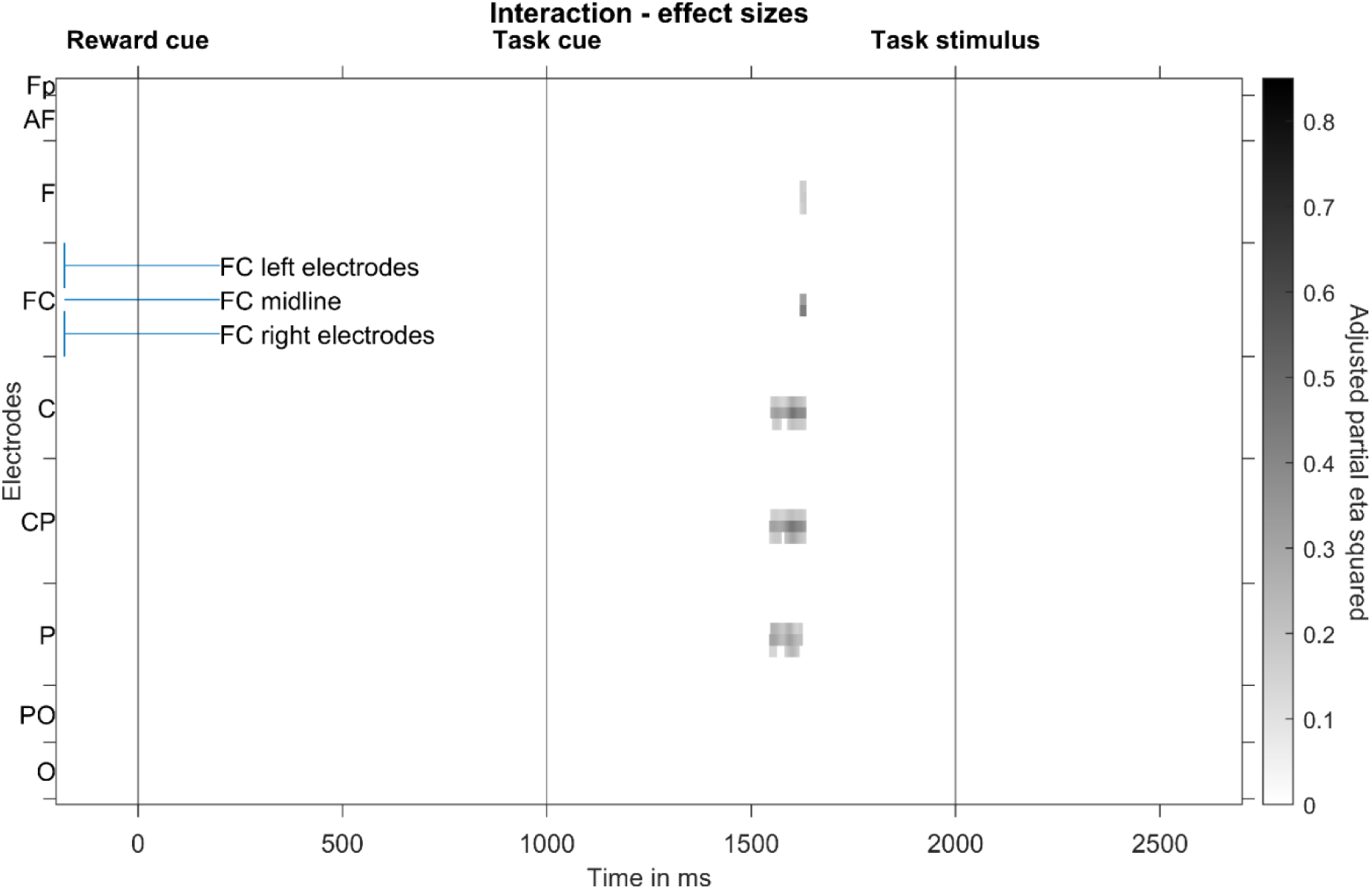
Result of cluster-based permutation analysis testing for the interaction between preciseness and reward. The analysis resulted in one significant cluster. Effect sizes (adjusted partial eta squared) at significant time-electrode combinations of this cluster are shown. Electrodes are lined up along the y axis, ranging from frontal to occipital rows, whereas each row’s electrodes are lined up from left to right (see labelling of FC electrode row).

Figure 10 shows the corresponding ERPs with topographic maps. As can be seen in the figure, in the interaction time interval, precisely cued bonus trials show a stronger negativity at central electrodes than all other conditions.

**Figure 10.**
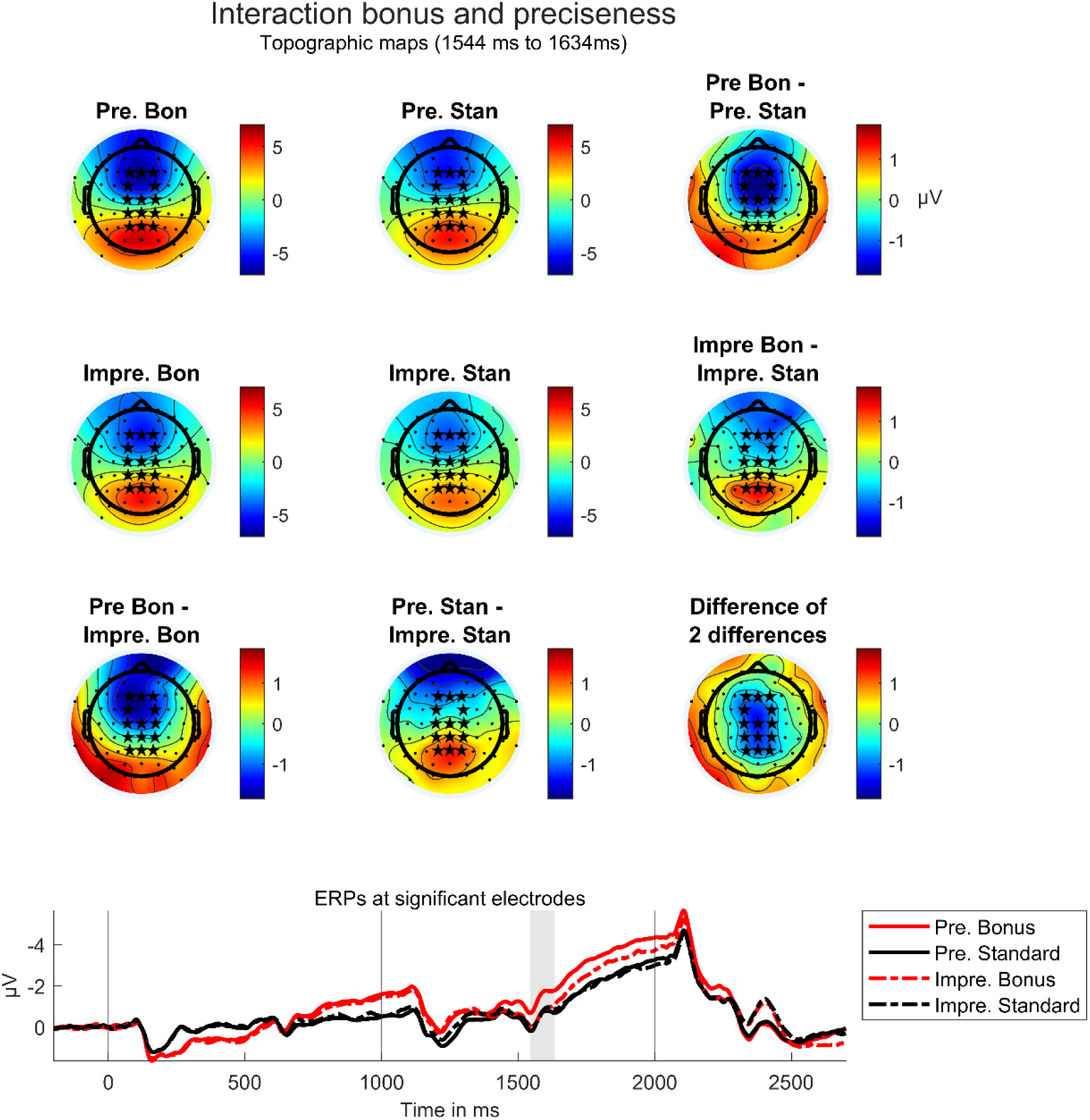
Significant interaction between task cue preciseness and reward according to a cluster-based permutation test. Top: Topographic maps for time intervals in which a significant interaction was found. Asterisks mark electrode at which the effect is significant. Depicted are single condition topographies (quartet at the top-left corner) and condition differences (outer right line and bottom line). The difference of the bonus effect in precisely cued trials minus the bonus effect in imprecisely cued trials (difference of differences, see right-bottom corner) is the same than the difference between the preciseness effect in bonus trials and the preciseness effect in standard trials. It shows that the interaction is strongest at central electrodes. Pre.: Precise; Impre.: Imprecise; Bon: Bonus; Stan: Standard. Bottom: ERPs averaged over significant electrodes. The grey patch marks the time interval at which the effect is significant, hence 1544 ms to 1634 ms after reward cue onset).

## Discussion

In this study, we examined how an increase of cognitive control due to an external factor, namely performance contingent reward prospects, can target more vs. less specific processes. In a cued task switch ERP study, we manipulated both, the availability of a monetary reward in case of good performance, and the preciseness of task-relevant information given prior to the task. We hypothesized that reward would affect task preparation in a task-specific manner and that the reward effect would flexibly adapt to situational factors that determine how specific the reward effect can be.

We can first state that our intended experimental manipulation was successful. As in earlier studies (Mazaheri et al., 2014; Sokoliuk et al., 2019), we obtained strong behavioral effects of the factor preciseness, both regarding accuracy and response times. This indicates that participants indeed made effective use of the additional pre-task information in precisely cued trials. Hence, preparatory processes in precisely cued trials reflect more specific task preparation than anticipatory processes in imprecisely cued trials.

This brings us to the question to what extent *reward* modulated specific and unspecific processes. As expected (Botvinick & Braver, 2015), we observed a significant main effect of reward in behavioral data, revealing a better performance when participants anticipated a reward than when they did not. A significant difference was only found for response times and not for response accuracies, a pattern which has earlier been observed when manipulating performance contingent reward in classification tasks (Kleinsorge & Rinkenauer, 2012; Liegel et al., 2022). Both in precisely and in imprecisely cued trials, participants performed better in bonus than in standard trials (compare Figure 2). This result was mirrored in EEG data in preparatory time intervals: We obtained significant ERP amplitude modulations by reward expectation, both in precisely and imprecisely cued trials, regarding an ERP slow wave (CNV) which appeared prior to the task stimulus. More specifically, in line with previous research on performance contingent reward prospects, the amplitude of the fronto-central CNV was more negative in rewarded than in unrewarded trials and this modulation was more frontal in earlier time intervals and then became more central (Schevernels et al., 2014; van den Berg et al., 2014). We can conclude so far that reward prospect did affect preparatory processes in precisely and imprecisely cued trials.

Since specific task preparation is not possible in imprecisely cued trials, it can be concluded that the observed effects of the reward manipulation reflect a modulation of imprecise processing. Regarding precise task cues, however, one might ask if reward really affected the specific preparatory processes that – as the behavioral data suggests – took place. The CNV has been shown to reflect both motor and attentional preparation to a task and might reflect cortical excitability in sensory-motor areas (Brunia et al., 2011; Kononowicz & Penney, 2016). Importantly, it is thought to reflect task-specific processes. More exactly, CNV amplitudes have been shown to reflect activity in multiple brain areas and typically show a task-specific distribution (Brunia et al., 2011; Kononowicz & Penney, 2016). In addition, the nature of an upcoming task can be predicted from EEG measures in the CNV time and frequency range (Hoxha et al., 2023). Our finding of an influence of reward on CNV amplitude, in both precisely cued and imprecisely cued trials, hence strongly suggests that reward modulated specific anticipatory top-down attention in precisely cued trials and unspecific top-down attention in imprecisely cued trials.

This assumption is strongly supported by another result regarding preparatory EEG activity: During an early CNV interval, we found an interaction between the factors preciseness and reward at central electrodes. A more pronounced negativity arose for the precisely cued bonus condition than for all other conditions. The effect is topographically and time-wise located at the transition between task cue P3 and early CNV. In addition, a trend for a stronger reward effect in precisely than imprecisely cued trials can be observed throughout the whole CNV time interval, although not significant with the – compared to other ERP studies – rather conservative multiple comparison correction applied here. It is plausible that it is the earliest time interval within the CNV component, hence the transition from a P3b to a CNV processing stage that is affected: The P3b is thought to reflect stimulus categorization and working memory updating (Glazer et al., 2018), while the early CNV reflects the selection of a preparatory strategy based on memory and task cue characteristics (Brunia et al., 2011). The result pattern suggests that the initiation of preparatory processes starts earlier and/or is more efficient in precisely cued bonus trials than in all other trial types, resulting in an especially strong preparation throughout the whole CNV time interval for precisely cued bonus trials.

To conclude, the reward manipulation in our study influenced both, precise and imprecise task preparation and both was affected differently. This is well in line with our hypothesis according to which reward affects task-specific top-down attention and flexibly adapts to different levels of task specification determined by situational factors.

Besides the adaption of reward effects over different task preparation situations, we were also interested in the adaption of reward effects over time. We aimed to understand if reward effects already occur at time intervals in which tasks are ill defined and how reward effects change as a result of a better task specification. Previous research suggests that the expectation of a reward is related to a stronger utilization of pre-stimulus information (Locke & Braver, 2008) which is associated with an increase of both preparatory and sustained attentional control (Hall-McMaster et al., 2019; Jimura et al., 2010; Sawaki et al., 2015). These results have been interpreted to indicate an increase of proactive control in case of reward prospects (Botvinick & Braver, 2015). We here want to extend these findings by examining variations in time regarding the specificity of the reward influence on proactive control. We used a task cue which divides the preparatory time interval in a phase during which only general task preparation was possible and another phase during which also specific task preparation took place. An important aspect of our experimental design in this regard is that one quarter of all trials were break trials. This means that prior to the task cue, participants did not even know whether a task will be presented or not. We were interested if the well-known influence of reward on proactive control generalizes to preparatory activity in such phases of high uncertainty represented here by the time interval prior to the task cue.

We found indeed strong evidence that reward modulated preparatory activity prior to the task cue. A more pronounced fronto-central negativity could be observed in bonus trials than in standard trials. We refer to this as stimulus-preceding negativity (SPN) in line with Brunia and colleagues (2011). Central slow waves prior to cues with instructional content have been observed before. They have been interpreted to reflect anticipatory attention (van Boxtel & Böcker, 2004). Much more research has been done on the SPN prior to feedback stimuli. This line of research also strongly suggests that negative going slow waves reflect anticipatory attention to the subsequent stimulus (Brunia et al., 2011). On the one hand, the modulation of SPN amplitude by reward could hence indicate that participants allocated more attentional resources to task cue processing in bonus than in standard trials. An alternative explanation is that the CNV prior to target onset starts already prior to the task cue, is then superimposed by task cue evoked potentials and continues after task cue processing. The modulation of pre-task cue slow waves would then represent an effect in a very early CNV. In line with that, a very early start of reward effects on slow wave amplitudes has been observed earlier (Pornpattananangkul & Nusslock, 2015). This would rather indicate that the observed effect in the SPN time interval reflects preparation to *task* processing rather than to *task cue* processing. Either way, results strongly indicate that there are reward-related differences in very early preparatory attentional top-down control. Note that we consider these early effects to be unspecific as no specific information has yet been provided at this time interval and any preparation that occurs must include all possible options that might or might not follow.

We also obtained a significant correlation between the pre-task cue bonus effect in the EEG and the bonus effect in response times. This indicates that the reward effect reflected by SPN amplitude modulation is functionally relevant for target processing. To conclude, we here observe an early and unspecific, functionally relevant reward effect that changes to a more specific one following precise task cues. Influences of reward on task preparation can thus flexibly adapt when information from the environment is integrated into current goals. It might change over time which top-down processes are influenced by reward prospects, with both more general and more specific processes being potential targets.

Results so far support our hypothesis that the effect of performance-contingent reward is as task-specific as the current situation allows it be, which is accompanied by the finding that the reward effect is highly flexible regarding both adaptations over time and adaptations across instructional contents. Interestingly, behavioral results additionally indicated that more general and more specific reward effects do not only flexibly differ regarding their underlying neural correlates but also regarding behavioral outcomes: We obtained a significant interaction in response times between task cue preciseness and reward, with stronger reward effects occurring in precisely cued than in imprecisely cued trials. This result suggests that task specification and reward are factors that influence each other or, more precisely, strengthen each other. The effect is especially interesting in combination with the EEG results: According to the EEG data, reward exerts its influence on performance mainly during the preparatory processing stage. We find, amongst others, a significant effect of reward on the CNV amplitude prior to the task stimulus, as well as an interaction regarding this ERP component which mirrors the behavioral interaction effect. In contrast, there is not even a trend in the data indicating that stronger *post*-target reward influences occur in imprecisely cued trials than in precisely cued trials to compensate for the worse preparation. Thus, we can conclude that effects of reward on *task preparation* are behaviorally more effective under precise than under imprecise task specification.

### Limitations and implications for future research

The significant behavioral interaction between task cue preciseness and reward might be a starting point for more applied research: If the interplay of task specification and reward can be replicated and transferred to more applied settings, it might guide the implementation of reward and pre-task information in different areas from teaching to security management in industrial production. In a first step, it would be interesting to understand if the effect is restricted to situations in which processing speed is rewarded or otherwise relevant. The fact that an increase of proactive control under performance contingent reward has been observed not only for response time but also for working memory tasks (Jimura et al., 2010) increases the likelihood that the interaction effect observed here might also generalize to non-response time tasks.

Besides the behavioral interaction effect, our results in its entirety open up opportunities for future research. This study provides a first step towards the understanding in how far changes in attentional control which are triggered by reward prospects concern more general or more specific attentional selection processes, and how this depends on situational factors. Our results do not allow, however, to draw conclusions about the exact nature of specific and general effects. When we learn new categories and generalized concepts, we create representations in the brain which are more than just the sum of the representations of single category exemplars (Freedman et al., 2001; Freedman & Assad, 2016; Freedman & Miller, 2008; Mansouri et al., 2020). Generalized concept representations contain typical category features but no exemplar-specific details so that the generalized representation fits to all exemplars of the category. Hence, a generalized task goal representation should contain characteristic information about all possible tasks, but no task-specific details. We do not know, if in imprecisely cued trials in this study, such generalized task goal representations were created and if reward affected such generalized representations and related selection processes. It has been found earlier that reward prospects *can* affect such generalized processes (Liegel et al., 2022). Future research might ask under which circumstances reward affects what type of specific and unspecific attentional selection processes.

### Conclusion

In the present study, we found effects of performance contingent reward on task performance and on preparatory attentional control, reflected in ERP slow wave amplitudes, for both specific and unspecific task specification. The specific reward effect, that is, the effect of reward on specific task goals and related top-down attention, differed from the unspecific effect – both in behavioral and EEG data. The pattern of results indicates that in the case of a specific task cue, reward affects different, namely more specific, processes than in the case of an unspecific task cue and that the specificity of the reward effect flexibly adapts to current task requirements. Moreover, general reward effects seem to be flexible enough to become more specific over time. Finally, our results suggest that specific reward effects are behaviorally more effective than general ones.

First, these results are interesting in the context of performance contingent reward. Previous research suggests that reward prospects can increase attentional resource allocation (Botvinick & Braver, 2015; Krebs & Woldorff, 2017). Here, we strongly confirm that assumption and extend it: Reward can target highly specific resource allocation processes. Which attentional processes are enhanced depends on the situation and this choice of processes is flexible enough to change over time when new information is provided. Behavioral performance benefits more from a case where reward affects specific processes than from a case where reward affects unspecific processes. Performance-contingent reward can thus result in more vs. less specific brain processes associated with different behavioral outcomes. There is no unique neural signature of reward prospects and it is worth to differentiate between more specific and more general effects.

Second, on a broader sense, results are interesting in the context of increases of attentional control in general. Results confirm earlier findings in the context of performance monitoring: An increase of attentional control varies in its specificity depending on the exact situation (Braem et al., 2014). There seem to be different types of environmental influencing factors triggering an increase in cognitive control: On the one hand, factors such as conflict adaptation trigger an influence on attentional control that induce more specific changes, on the other hand, factors such as error processing trigger more general changes (Forster & Cho, 2014; Ullsperger, 2017). Our results suggest that performance contingent reward prospects triggers task-specific changes, or more exactly changes which are as task-specific as they can be according to the given task specification. In this regard, the present study adds to a better understanding how environmental factors differ regarding their influence on attentional selection processes.

To sum up, results highlight how flexibly the cognitive system can adapt to its environment: Not only the content of goals and the strength of attentional control but also the specificity of goal updates can be flexibly modified in line with current requirements.

## Data / code availability statement

The datasets analyzed during the current study and the analysis code will be available in the OSF Repository, https://osf.io/n26u9/, upon publication.

## Acknowledgements

We would like to thank Pia Deltenre and Karin Lukaszewski for their assistance in collecting the data as well as Tobias Blanke for programming the experiment.

## Additional Information

Portions of these findings were presented as a poster at the CNS Annual Meeting, San Francisco, 2023. We have no conflicts of interest to disclose.

## Author contributions

N.L..: Conceptualization; Project administration; Investigation; Data curation; Formal analysis; Visualization; Writing-original draft; Writing-review & editing. D.S.: Conceptualization; Writing-review & editing; Supervision. E.W.: Writing-review & editing; Resources; Supervision. L.K.: Writing-review & editing; Supervision. S.A.: Conceptualization; Project administration; Writing-review & editing; Resources; Supervision.

